# Accessible, curated metagenomic data through ExperimentHub

**DOI:** 10.1101/103085

**Authors:** Edoardo Pasolli, Lucas Schiffer, Paolo Manghi, Audrey Renson, Valerie Obenchain, Duy Tin Truong, Francesco Beghini, Faizan Malik, Marcel Ramos, Jennifer B. Dowd, Curtis Huttenhower, Martin Morgan, Nicola Segata, Levi Waldron

## Abstract

We present curatedMetagenomicData, a Bioconductor and command-line interface to thousands of metagenomic profiles from the Human Microbiome Project and other publicly available datasets, and ExperimentHub, a platform for convenient cloud-based distribution of data to the R desktop. The resource provides standardized per-participant metadata linked to bacterial, fungal, archaeal, and viral taxonomic abundances, as well as quantitative metabolic functional profiles. The datasets can be immediately analyzed in R or other software with a minimum of bioinformatic expertise and no preprocessing of data. We demonstrate identification of taxonomic/functional correlations, an investigation of gut “enterotypes”, and a comparison of the accuracy of disease classification from different data types. These documented analyses can be reproduced efficiently on a laptop, without the barriers of working with large-scale, raw sequencing data. The building and expansion of curatedMetagenomicData is based entirely on open source software and pipelines, to facilitate the addition of new microbiome datasets and methods.

---

The human microbiome has emerged as a key aspect of human biology and has been implicated in many etiologies. Shotgun metagenomic sequencing is the most high-resolution approach available to study taxonomic composition and functional potential of the human microbiome, and an increasing amount of published data are available for re-use. These public data resources allow the possibility of rapid, inexpensive hypothesis testing for specific diseases and environmental niches, and meta-analysis across multiple related studies. However, several factors prevent the research community from taking full advantage of these public resources. Barriers include the substantial investments of time, computational resources, and specialized bioinformatic expertise required to convert them to analyzable form, and inconsistencies in annotation and formatting between individual studies.

To overcome these challenges, we developed the curatedMetagenomicData data package (described at https://waldronlab.github.io/curatedMetagenomicData/) for distribution through the Bioconductor^1^ ExperimentHub platform (see **Supplementary Methods**). curatedMetagenomicData provides highly curated and uniformly processed human microbiome data including bacterial, fungal, archaeal, and viral taxonomic abundances, in addition to quantitative metabolic functional profiles and standardized per-participant metadata. Data resources are accessible with a minimum of bioinformatic knowledge, while integration with the R/Bioconductor environment allows full flexibility for biologists, clinicians, epidemiologists, or statisticians to perform novel analyses and methodological development. We produced these resources by (i) downloading the raw sequencing data, (ii) processing it through the MetaPhlAn2^2^ and HUMAnN2^3^ pipelines, (iii) manually curating sample and study information, (iv) creating a pipeline to document and represent the above results as integrative Bioconductor objects, and (v) working with the Bioconductor core team to develop the ExperimentHub platform for efficient distribution. ExperimentHub is a novel platform for scalable distribution of experimental data to the R desktop. It allows distributers and downloaders to interact through a standard R software package, with documented convenience functions for accessing metadata through a Bioconductor-hosted SQL database, and bulk data through Amazon S3 buckets. Users of the data can browse per-dataset documentation and metadata, then download any dataset, along with curated patient and specimen information, directly into R or from the command line with a single operation. To date (development version 1.3.7), we have packaged samples from multiple body sites profiled by the Human Microbiome Project^4^ and from 25 other large metagenomic studies. These total 5,716 samples, spanning 34 diseases and 28 countries. The full pipeline is summarized in **Figure 1**, and datasets are listed in **Supplemental Table 1**.

**Figure 1:**
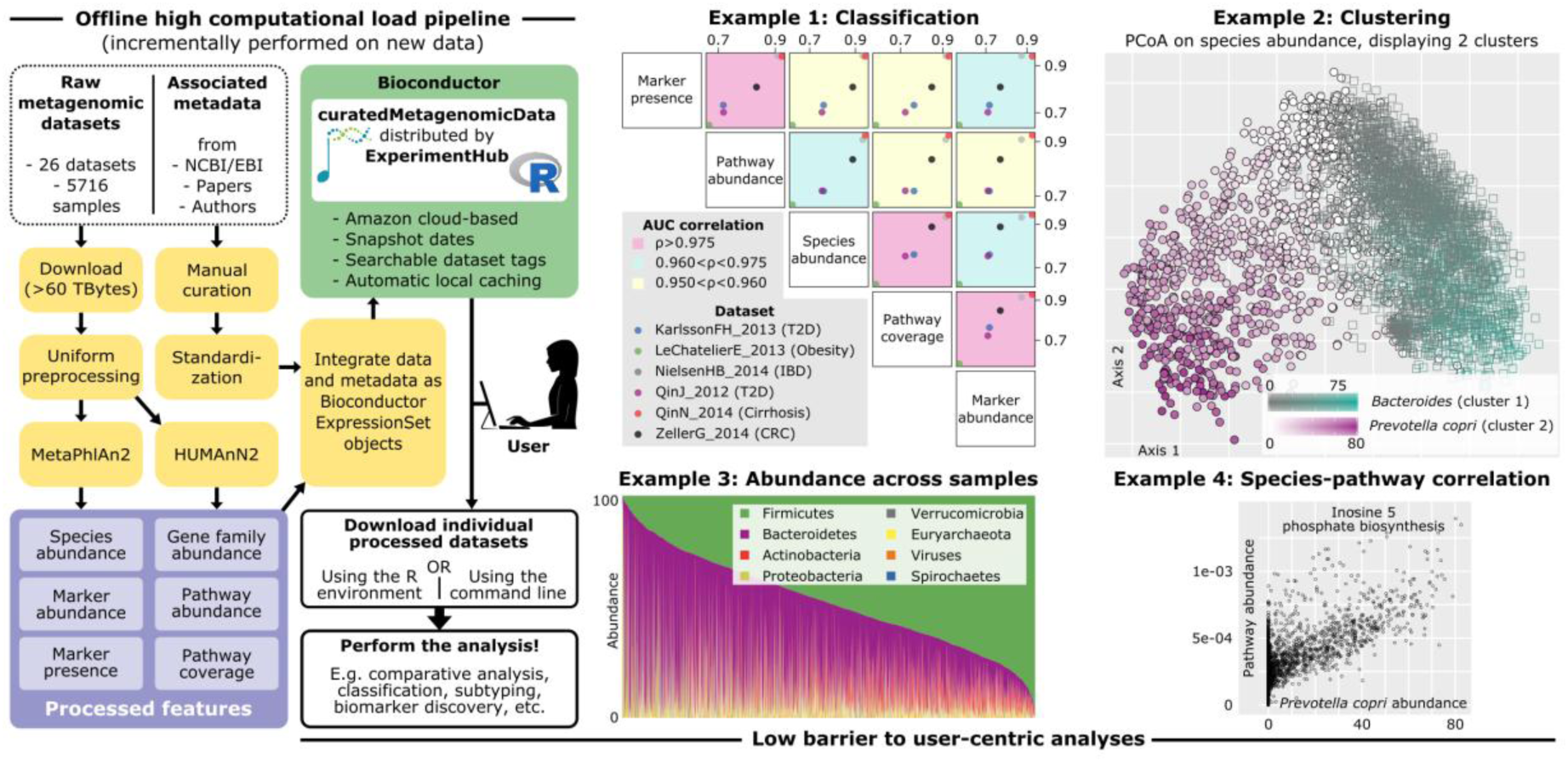
curatedMetagenomic Data production pipeline and examples of enabled analyses. The high computational load pipeline (left) processes raw metagenomic sequence data to produce taxonomic and functional profiles, integrates these with curated sample data, then documents and packages these for distribution through ExperimentHub as the curatedMetagenomicData package. Example 1: Six different classification problems of health status were attempted using a random forest algorithm and cross-validation to estimate prediction accuracy. The classification problems range from easy (AUC > 0.9) to harder (AUC < 0.7), but five different data products (three taxonomic and two functional) provide nearly identical performance on each classification problem. Example 2: Unsupervised clustering of human gut samples shows two weakly separated clusters, one characterized by *Bacteroides* prevalence and the other characterized by *Prevotella copri* prevalence. Example 3: At the phylum level, the human gut microbiome is characterized primarily by a Firmicutes/Bacteroidetes gradient, with loads of other bacterial phyla, archaea, and viruses varying from negligible to over 50%. Example 4: *Prevotella copri* and inosine 5 phosphate biosynthesis are the most correlated species-pathway pair, suggesting functional difference along the *Prevotella copri* gradient shown in Example 2. A heatmap of top species-pathway pairs is provided as **Supplemental Figure 4**. These analyses are performed using the script provided in the vignettes/extras package directory.

We performed several analyses that are made much more straightforward and powerful by curatedMetagenomicData and the statistical, visualization, and microbial ecology tools available in R/Bioconductor. Using a random forests algorithm we used three different taxonomic data types (species abundance, genetic marker presence and abundance) and two functional abundance profiles (pathway abundance and coverage), to develop predictive models of diabetes, inflammatory bowel disease, cirrhosis, colorectal cancer, and obesity. Cross-validation prediction accuracy varied substantially for these different applications, but in all cases the five data types provided nearly identical accuracy (Figure 1 example 1). Second, we performed unsupervised clustering of human gut microbiome profiles. In a large combined dataset (n=3667), we observed that microbial communities are strongly patterned by abundance of *Prevotella copri* and *Bacteroides spp* (Figure 1 example 2), consistent with the analysis of Koren *et al.*^5^, but not with the three-enterotypes hypothesis of Arumugam *et al.*^6^. Third, we visualized the continuum of the Firmicutes/Bacteroidetes gradient in gut microbiomes as reported previously^4^, but these abundances can now be investigated for thousands of microbial species (Figure 1 example 3). Finally, we ranked all taxa/pathway pairs by magnitude of correlation in samples. The highest-correlation pair shown demonstrates a strong relationship between *Prevotella copri* abundance and inosine 5 phosphate biosynthesis (Figure 1 example 4), suggesting functional differences along the gradient shown in example 2. These and other analyses (**Supplemental Figures 1-5)**, would be very large undertakings using less curated databases such as IMG/M or EBI Metagenomics, but are straightforward, documented, and reproducible analyses using curatedMetagenomicData.

We present the first curated integration of large-scale metagenomic data and make it readily usable by broad scientific communities. With overall and per-dataset documentation, and integration with R/Bioconductor, curatedMetagenomicData enables efficient hypothesis testing and development of statistical methodologies specifically for microbiome data. The automated pipeline developed here will enable continued expansion of the resource by the current team and contributing members of the community, as described in the supplementary *Package maintenance.* By allowing researchers to bring their own expertise to the analysis of metagenomic data without the need for extensive bioinformatic expertise, curatedMetagenomicData greatly expands the accessibility of public data for study of the human microbiome.

## Methods

### Available datasets

To date (development version 1.3.7), we packaged a total of 5,716 publicly available shotgun metagenomic samples coming from 26 large-scale studies (see **Supplemental Table 1**). All these metagenomes have been sequenced on the Illumina platform at an average depth of 45 M reads.

Twelve of these studies were performed to assess the association of the human gut microbiome with different diseases. In particular, four studies were devoted to the characterization of the human microbiome in colorectal cancer patients: FengQ_2015^11^ (154 samples, 93 cases), VogtmannE_2016^27^ (110 samples, 52 cases), YuJ_2015^29^ (128 samples, 75 cases), and ZellerG_2014^30^ (199 samples, 133 cases). Heitz-BuschartA_2016^12^ includes a total of 53 samples, 27 of which are associated with type 1 diabetes (T1D). KarlssonFH_2013^13^ sampled on European women and includes 53 type 2 diabetes (T2D) patients, 49 impaired glucose tolerance individuals and 43 normal glucose tolerance individuals. QinJ_2012^20^ sampled an additional T2D dataset and is composed by 170 Chinese T2D patients and 193 non-diabetic controls. LeChatelierE_2013^14^ includes 123 non-obese and 169 obese individuals. LomanNJ_2013^16^ includes 43 samples from patients with life-threatening diarrhea during the 2011 outbreak of Shiga-toxigenic Escherichia coli (STEC) O104:H4 in Germany. NielsenHB_2014^17^ focuses on inflammatory bowel disease (IBD) and comprises a total of 396 samples, 21 of which are from Crohn’s disease patients and 127 from ulcerative colitis patients. QinN_2014^21^ includes 123 patients affected by liver cirrhosis and 114 healthy controls. VincentC_2016^26^ focused on microbiota dynamics in response to hospital exposures and Clostridium difficile colonization infection in a total of 229 samples.

We included also four datasets that investigated gut configuration in hunter-gatherer or non-westernized populations. BritoIL_20168 considered agrarian Fiji islanders for a total of 312 samples, including also some samples from the oral cavity. LiuW_2016^15^ investigated 110 Mongolian adults. Obregon-TitoAJ_2015^18^ sequenced 58 samples, which include hunter-gatherer and traditional agriculturalist communities in Peru. RampelliS_2015^22^ comprises 38 samples, part of which were collected from Hadza hunter-gatherers of Tanzania.

Additional datasets were acquired entirely from healthy subjects. AsnicarF_20177 collected 24 samples for studying vertical microbiome transmission from mothers to infants. RaymondF_2016^23^ acquired 72 samples to evaluate effects of a standard antibiotic treatment on the microbiome. SchirmerM_2016^24^ investigated 471 samples to link the microbiome to inflammatory cytokine production capacity. VatanenT_2016^25^ considered 222 infants in Northern Europe from birth until age three for a total of 785 samples. XieH_2016^28^ investigated 250 adult twins to evaluate genetic and environmental impacts on the microbiome.

Some datasets not strictly related to the gut microbiome are also taken into account. Castro-NallarE_20159 collected 32 samples from the oral cavity to investigate the oropharyngeal microbiome in individuals with schizophrenia. HMP4 includes 749 samples collected for the Human Microbiome Project from five major body sites (i.e., gastrointestinal tract, nasal cavity, oral cavity, skin, and urogenital tract). Finally, three datasets focused on the skin microbiome. OhJ_2014^19^ is composed by 291 samples collected from several different skin sites in healthy conditions. Skin samples but from patients affected by atopic dermatitis and psoriasis were acquired in ChngKR_2016^10^ (78 samples) and TettAJ_2016 (97 samples, BioProject accession number PRJNA281366), respectively.

### Raw data pre-processing

Approximately 63 TB of raw sequencing data were downloaded from public repositories. All samples were subject to standard pre-processing as described in the SOP of the Human Microbiome Project^4^, without however the step of human DNA removal as these publicly available metagenomes were deposited free of reads from human DNA contamination.

### MetaPhlAn2 profiling and data products

MetaPhlAn22 (v2.0) was ran on the pre-processed reads with default parameters to generate microbial community profiles (from kingdom- to species-level) including Bacteria, Archaea, microbial Eukaryotes and Viruses. These profiles were generated from ^~^1 M unique clade-specific marker genes identified from ^~^17,000 reference genomes (^~^13,500 bacterial and archaeal, ^~^3,500 viral, and ^~^110 eukaryotic). MetaPhlAn2 has the capability of characterizing organisms at a finer resolution using non-aggregated marker information (“-t marker_pres_table” and “-t marker_ab_table” mode). Single marker-level profiles were then merged in samples versus markers tables removing markers there were never detected in any samples.

Such processing resulted in three data products: i) species-level relative abundance (denoted as “metaphlan_bugs_list” in the package); ii) marker presence (“marker_presence”); and iii) marker abundance (“marker_ abundance”). Species abundance is expressed in percentage and sum up to hundred within each sample when selecting a single taxonomic level. Marker presence and marker abundance assume binary and real values, respectively.

### HUMAnN2 profiling and data products

HUMAnN23 (v0.7.1) was run on the pre-processed reads with default parameters for profiling the presence/absence and abundance of microbial pathways in the community. The mapping was done using the full UniRef90 database (~11 GB), which enabled identifying also protein families without functional annotations. Three main outputs were generated: gene family abundance, pathway abundance, and pathway coverage. The two abundance output files were normalized in terms of relative abundance through the “humann2_renorm_table” (“--units relab” mode).

In this way, three additional data products were produced: i) normalized gene family abundance (denoted as “genefamilies_relab” in the package); ii) normalized pathway abundance (“pathabundance_relab”); and iii) pathway coverage (“pathcoverage”). Features assume values in the range [0, 1], where the two normalized abundance profiles sum up to 1 when excluding species-specific contributions.

### Creation of curatedMetagenomicData

To create the curatedMetagenomicData package, processed data, in the form of tab-delimited files, from the MetaPhlAn2 and HUMAnN2 pipelines and patient-level metadata are compressed into a single archive file per dataset. Then from within the R/Bioconductor environment a single function is used to process the compressed archive, create documentation, and add to curatedMetagenomicData, with internal intermediate steps as follows. First, patient-specific metadata is read in using the readr package (https://CRAN.R-project.org/package=readr), filtered using the dplyr (https://CRAN.R-project.org/package=dplyr) and magrittr (https://CRAN.R-project.org/package=magrittr) packages, and coerced to the appropriate format. Study-level metadata is then created by querying PubMed using the RISmed package (https://CRAN.R-project.org/package=RISmed), which collects citation information of published studies that can then be coerced to the appropriate format. Finally, patient-level sample data is read in (again using the readr package), merged, standardized, and used to create Bioconductor ExpressionSet objects^34^ featuring the patient and study-level metadata. Within each study, processed data is separated into six data products, as highlighted above, and further separated by bodysite so as to allow for efficient search and data transfer.

Once data from the MetaPhlAn2 and HUMAnN2 pipelines have been processed into Bioconductor ExpressionSet objects, documentation, package metadata, and upload to ExperimentHub are accomplished using developer functions available in curatedMetagenomicData. Documentation is automatically produced from the ExpressionSet objects using roxygen2 (https://CRAN.R-project.org/package=roxygen2), although this may change in the future. Package metadata is also produced from the ExpressionSet objects and used in the creation of ExperimentHub records, with further details concerning ExperimentHub below. Finally, a convenience function is provided to write a shell script to upload all data to ExperimentHub, such that the error-prone process of working with Amazon Web Services (AWS) Command Line Interface (CLI) is trivial.

### Bioconductor object classes

*curatedMetagenomicData* data objects are represented using the Bioconductor ExpressionSet S4 class^34^. This class links numeric microbiome data with subject information and whole-experiment level data, while maintaining correct alignment between numeric microbiome data subject data during subset operations. The following ExpressionSet slots are populated in each data product:

- *experimentData*: “MIAME” class object providing study-level information - Pubmed ID, authors, title, abstract, sequencing technology, etc. Extracted using experimentData(object).
- *phenoData*: “AnnotatedDataFrame” class object providing specimen-level information - subject IDs, disease, body site, number of reads, etc. Extracted using pData(object) or phenoData(object).
- *assayData*: matrix class object providing taxonomic or pathway abundances. Extracted using exprs(object).

ExpressionSet objects can be analyzed for differential abundance using popular Bioconductor packages for RNA-seq such as *limma, edgeR, and DESeq2.* For MetaPhlAn2 abundances, however, it is more convenient to convert these to *phyloseq* objects for analysis with the *phyloseq* Bioconductor package for phylogenetics, using the *ExpressionSet2phyloseq* function from *curatedMetagenomicData*. Phyloseq objects additionally represent taxonomy and phylogenetic distances, and enable straightforward calculation of alpha and beta diversity measures, ordination plots, and other phylogenetic-specific analyses.

### ExperimentHub

*curatedMetagenomicData* datasets are distributed through ExperimentHub, a new Bioconductor software package we developed to provide programmatic access to experimental data files stored in the Amazon Web Services (AWS) cloud. All data (referred to as “resources”) in ExperimentHub have undergone some level of curation and are provided as R/Bioconductor data structures instead of in raw format. Data sets are generally a collation of different sources combined by disease or cohort or data used in a published experiment or short courses.

The two primary components of ExperimentHub are the data files and the metadata describing them. Files are stored in AWS S3 buckets and the metadata in a database on the ExperimentHub server. The database version is reflected in the “snapshot date” which is updated whenever the database is modified. Users interacting with ExperimentHub can select a specific snapshot date which, along with the version of R / Bioconductor, modifies which resources are exposed.

ExperimentHub resources are accessed by invoking ExperimentHub() to create an ‘ExperimentHub’ object, e.g., hub <- ExperimentHub(). This call downloads the database of metadata from the ExperimentHub server and caches it locally. The ‘hub’ of metadata can be searched with the query() function and subset by numerical index or ‘EH’ identifier. Once a resource is identified, the double-bracket method (‘[[’) will initiate the download. Downloaded resources are cached locally enabling fast repeated access to the data. When a resource is loaded in an R session, the accompanying software package is also loaded ensuring all documentation and helper functions are readily available. A second option for accessing the data is to invoke the resource name as a function, e.g., data123(). In this approach, the creation and searching of the ‘hub’ is not exposed to the user and does not require knowledge of ExperimentHub objects.

Resources are added to ExperimentHub by creating a software package according to the guidelines in the ExperimentHubData vignette (https://bioconductor.org/packages/release/bioc/vignettes/ExperimentHubData/inst/doc/ExperimentHubData.html). The software package includes man pages and a vignette documenting expected use as well as functions to create the resource metadata. If desired, the author may include additional functions for resource discovery and manipulation. Data are stored separately in AWS and are not part of the software package; this separation enables lightweight installation of the package regardless of the size of the data.

### Accessing curatedMetagenomicData objects in R

Within the R/Bioconductor environment there are two distinct methods for accessing data, depending on the needs of the end-user. In the case that a specific dataset is desired and its name is known, then convenience functions have been provided for all datasets and calling the function will retrieve the dataset from ExperimentHub. Otherwise, if no specific dataset is desired, it is possible to search through all datasets and return those matching a pattern (e.g., all datasets from the stool bodysite). This method also features wildcard search to allow for powerful selection and can return either a list of references to the datasets or download the datasets from ExperimentHub. The later search method is of particular use in conducting cross validation studies using curatedMetagenomicData, as it provides for highly specific filtering conditions.

### Accessing curatedMetagenomicData from the command line

A convenience command-line interface is provided for users who do not want to use the R or Bioconductor framework for the analysis. The command-line program is invoked with the names of one or more datasets with optional wildcard expansion, and provides flags for including specimen information in addition to microbiome data, and for returning relative abundances or counts. Datasets are written to disk as tab-separated value plain text files.

### Examples of enabled downstream tasks: supervised classification analysis

We considered six different classification problems of health status to evaluate capabilities of disease classification from gut microbial profiling (see Example 1 of **Figure 1** and **Supplemental Figure 1**). In KarlssonFH_2013, we discriminated between “healthy” and “T2D” subjects. We took into account 96 samples after excluding impaired glucose tolerance individuals. In LeChatlierE_2013, we discriminated between “lean” (BMI ≤ 25 kg m ^−2^) and “obese” (BMI ≥ 30 kg m ^−2^) subjects for a total of 265 samples. Individuals having an intermediate BMI (i.e., > 25 and < 30 kg m ^−2^) were excluded. NielsenHB_2014 was composed by a total of 396 samples, in which the “diseased” class included inflammatory bowel disease (IBD) patients affected by both “Crohn’s disease” and “ulcerative colitis”. In QinJ_2012 we considered a total of 344 samples and discriminated between “healthy” and “T2D” individuals. In QinN_2014, all the 237 available samples (subdivided into “healthy” and affected by “liver cirrhosis” subjects) were taken into account. Finally, in ZellerG_2014 we removed the individuals affected by “large adenoma”, which resulted in a total of 184 samples. “Cancer” patients were discriminated from “healthy” subjects, which included also persons affected by “small adenoma”.

We compared five different data products, three taxonomic (i.e., relative abundance, marker presence, and marker abundance) and two functional (i.e., normalized pathway abundance and pathway coverage). We subset relative abundance profiles to consider only species-level features, while the whole set of available features were taken into account for the other four data products.

The classification problems were attempted using the random forest algorithm through the R packages “randomForest” and “caret”. Original features were preprocessed (“preProc”) by centering (“center”), scaling (“scale”) and removal of zero-variance predictors (“zv”) procedures. Prediction accuracies were estimated using a 10-fold cross-validation approach (“method=repeatedcv” and “number=10” in the “trainControl” function). The two main parameters of the classifier were set in this way: i) the number of trees (“ntree”) was set to 500; ii) the number of variables randomly sampled as candidates at each split (“mtry”) were estimated through grid search. Area under the curve (AUC) values (**Figure 1**) were computed through the “auc” function in the R package “pROC”. The scatterplot matrix (**Figure 1**) was generated through the R package “gclus”, which provided possibility to i) rearrange the variables so that those with higher correlations are closer to the principal diagonal and ii) color the cells to reflect the value of the correlations. The “p ROC” package was also adopted to plot the receiver operating characteristic (ROC) curves (**Supplemental Figure 1**) using the “roc” function.

### Examples of enabled downstream tasks: unsupervised clustering analysis

To assess the presence of discrete clustering in the data (see Example 2 of **Figure 1** and **Supplemental Figure 3**), we merged taxonomic abundance data from all gut samples (excluding newborns), on which we calculated three distance measures using the R package “phyloseq”: the Bray-Curtis distance metric, the Jenson-Shannon divergence (JSD), and the square root of the Jenson-Shannon divergence (root-JSD). We then performed clustering against each of the three distance measures by partitioning around medoids using the R package “cluster”. We determined the optimal number of clusters based on the prediction strength (PS) using the R package “fpc”, and silhouette index (SI) using the R package “cluster”. We used a threshold of ≥ 0.90 for PS, and ≥ 0.75 for SI, to indicate strong clustering^5^. We additionally calculated the Calinski-Harabasz (CH) statistic for comparison to PS and SI, using the R package “fpc”.

### Package maintenance

We set up the curatedMetagenomicData to be scalable to the growing size of metagenomic datasets being produced and we plan to expand to over 10K total samples by the end of 2017, with dedicated personnel for the addition of processed metagenomic datasets. The curatedMetagenomicData pipeline directly uses output of the publicly available MetaPhlAn2 and HUMAnN2 packages, in a documented subdirectory structure for data “handoff” to our pipeline for incremental dataset addition to curatedMetagenomicData in ExperimentHub (https://github.com/waldronlab/curatedMetagenomicData/wiki).

Authors welcome the addition of new datasets provided they can be or already have been run through the MetaPhlAn2 and HUMAnN2 pipelines. Please contact the maintainer if you have a shotgun metagenomic dataset that would be of interest to the Bioconductor community.

### Availability and support

The curatedMetagenomicData package can be installed with a single command from an R installation with the current Bioconductor release or development version installed (BiocInstaller::biocLite(“curatedMetagenomicData”)). The package is described at https://waldronlab.github.io/curatedMetagenomicData/, including information on installation, datasets to be added in the near future, and example analyses. Requests for help should be raised at https://support.bioconductor.org with the tag *curatedMetagenomicData*. Bugs in code or curation should be reported using the issue tracker at https://github.com/waldronlab/curatedMetagenomicData/issues. Instructions for adding datasets or re-using parts or all of the pipeline for other purposes are provided on the wiki at https://github.com/waldronlab/curatedMetagenomicData/wiki.

### Reproducible analysis

All analyses presented in this manuscript are reproducible by the script PaperFigures.Rmd at https://github.com/waldronlab/curatedMetagenomicData/tree/master/vignettes/extras.

### Licensing

The curatedMetagenomicData package is licensed under the permissive Artistic 2.0 license.

## Acknowledgments

This work was made possible by the CUNY High Performance Computing Center, which is operated by the College of Staten Island and funded, in part, by grants from the City of New York, State of New York, CUNY Research Foundation, and National Science Foundation Grants CNS-0958379, CNS-0855217 and ACI 1126113. This work was supported in part by the European Union H2020 Marie-curie grant (707345) to E.P., the European Research Council (ERC-STG project MetaPG), MIUR “Futuro in Ricerca” RBFR13EWWI_001, the People Programme (Marie Curie Actions) of the European Union Seventh Framework Programme (FP7/2007-2013) under REA grant agreement no. PCIG13-GA-2013-618833, the LEO Pharma Foundation, and by Fondazione CARITRO fellowship Rif.Int.2013.0239 to N.S., the National Institute of Allergy and Infectious Diseases (1R21AI121784-01 to J.B.D. and L.W.) and the National Cancer Institute (1R03CA191447-01A1 and U24CA180996 to L.W.) of the National Institutes of Health.

## References

1. Huber, W. et al. Orchestrating high-throughput genomic analysis with Bioconductor. Nat. Methods 12, 115–121 (2015).

2. Truong, D. T. et al.et al.Nat. Methods 12,

3. Abubucker, S. et al. Metabolic reconstruction for metagenomic data and its application to the human microbiome. PLoS Comput. Biol. 8, e1002358 (2012).

4. Human Microbiome Project Consortium. Structure, function and diversity of the healthy human microbiome. Nature 486,

5. Koren, O. et al. A guide to enterotypes across the human body: meta-analysis of microbial community structures in human microbiome datasets. PLoS Comput. Biol. 9, e1002863 (2013).

6. Arumugam, M. et al.et al. Nature 473,

7. Asnicar, F. et al. Studying vertical microbiome transmission from mothers to infants by strain-level metagenomic profiling. mSystems 2, e00164–16 (2017).

8. Brito, I. L. et al. Mobile genes in the human microbiome are structured from global to individual scales. Nature 535, 435–439 (2016).

9. Castro-Nallar, E. et al. Composition, taxonomy and functional diversity of the oropharynx microbiome in individuals with schizophrenia and controls. PeerJ 3, e1140 (2016).

10. Chng, K. R. et al. Whole metagenome profiling reveals skin microbiome-dependent susceptibility to atopic dermatitis flare. Nat. Microbiol. 1, 16106 (2016).

11. Feng, Q. et al. Gut microbiome development along the colorectal adenoma-carcinoma sequence. Nat. Commun. 6, 6528 (2015).

12. Heintz-Buschart A. et al. Integrated multi-omics of the human gut microbiome in a case study of familial type 1 diabetes. Nat Microbiol. 2, 16180 (2016).

13. Karlsson, F. H. et al. Karlsson, F. H. et al. 498,

14. Le Chatelier, E. et al. Richness of human gut microbiome correlates with metabolic markers. Nature 500, 541–546 (2013).

15. Liu, W. et al. Unique features of ethnic mongolian gut microbiome revealed by metagenomic analysis. Sci. Rep. 6, (2016).

16. Loman, N. J. et al.et al.JAMA 309,

17. Nielsen, H. B. et al. Identification and assembly of genomes and genetic elements in complex metagenomic samples without using reference genomes. Nat. Biotechnol. 32, 822–828 (2014).

18. Obregon-Tito, A. J. et al.et al.Nat. Commun. 6, 6505 (2015).

19. Oh, J. et al.et al.Nature 514,

20. Qin, J. et al. Qin, J. et al. 490,

21. Qin, N. et al. Alterations of the human gut microbiome in liver cirrhosis. Nature 513,

22. Rampelli, S. et al. Metagenome sequencing of the Hadza hunter-gatherer gut microbiota. Curr. Biol. 25, 1682–1693 (2015).

23. Raymond, F. et al. The initial state of the human gut microbiome determines its reshaping by antibiotics. ISME J. 10, 707 (2016).

24. Schirmer, M. et al. Linking the human gut microbiome to inflammatory cytokine production capacity. Cell 167, 1125–1136 (2016).

25. Vatanen, T. et al. Variation in microbiome LPS immunogenicity contributes to autoimmunity in humans. Cell 165, 842–853 (2016).

26. Vincent, C. et al. Bloom and bust: intestinal microbiota dynamics in response to hospital exposures and Clostridium difficile colonization or infection. Microbiome 4, 12 (2016).

27. Vogtmann, E. et al. Colorectal cancer and the human gut microbiome: reproducibility with whole-genome shotgun sequencing. PLOS One 11, e0155362(2016).

28. Xie, H. et al. Shotgun metagenomics of 250 adult twins reveals genetic and environmental impacts on the gut microbiome. Cell Syst. 3, 572–584 (2016).

29. Yu, J. et al. Metagenomic analysis of faecal microbiome as a tool towards targeted non-invasive biomarkers for colorectal cancer. Gut 66, 7078 (2017).

30. Zeller, G. et al. Potential of fecal microbiota for early-stage detection of colorectal cancer. Mol. Syst. Biol. 10, 766 (2014).

31. Pasolli, E. et al. Machine learning meta-analysis of large metagenomics datasets: tools and biological insights. PLoS Comput. Biol. 12, e1004977 (2016).

32. Shi, D. et al. Structure and catalytic mechanism of a novel N-succinyl-L-ornithine transcarbamylase in arginine biosynthesis of Bacteroides fragilis. J. Biol. Chem. 281, 20623–20631 (2006).

33. Cusa, E. et al. Genetic analysis of a chromosomal region containing genes required for assimilation of allantoin nitrogen and linked glyoxylate metabolism in Escherichia coli. J. Bacteriol. 181, 7479–7484 (1999).

34. Falcon, S. et al. An introduction to Bioconductor’s ExpressionSet Class (2007).

